# Genome-wide characterization and identification of synonymous codon usage patterns in *Plasmodium knowlesi*

**DOI:** 10.1101/2021.01.01.425038

**Authors:** Manoj Kumar Yadav, Shivani Gajbhiye

**Affiliations:** Department of Biochemistry, Pt. J.N.M. Medical College, Raipur (CG) - 492001, India; Department of Bioinformatics, SRM University Delhi-NCR, Sonepat, Haryana, 131029, India

**Keywords:** Codon usage bias, Malaria, Mutation, translational selection pressure, Synonymous codons

## Abstract

Codon usage bias is a ubiquitous phenomenon occurring at both, interspecies and intraspecies level in different organisms. *P. knowlesi,* whose natural host is long-tailed Macaque monkeys, has recently started infecting humans as well. The genome as well as coding sequence data of *P. knowlesi* is used to understand their codon usage pattern in the light of other human infecting *Plasmodium* species: *P. vivax* and *P. falciparum.* The different codon usage indicators: GC content, relative synonymous codon usage, effective number of codon and codon adaptation index are studied to analyze codon usage in the *Plasmodium* species. The codon usage pattern is found to be less conserved in studied *Plasmodium* species, and changes species to species at the genus level. The codon usage pattern of *P. knowlesi* shows similarity to *P. vivax* as compared to *P. falciparum.* The ENC vs. GC3 study indicates that compositional constraints and translation selection is the decisive forces responsible for shaping their codon usage. The studies *Plasmodium* species shows a higher usage of A/T ending optimal codons. This favors the codon bias in *P. knowlesi* and *P. vivax* is due to high selection pressure and in *P. falciparum,* the compositional mutational pressure is a dominant force. In a nutshell, our finding suggests that the more or less similar codon usage pattern of *P. knowlesi* and *P. vivax* may suggest the similar host invasion and immune evasion strategies for disease establishment.

## Introduction

Malaria is an infectious disease caused by infected female mosquito-bite of the genus *Anopheles*. It is not a sexually transmitted disease but spread through organ transplantation, blood transfusion and also from mother to fetus. Malaria is caused by *Plasmodium* species, a parasitic protozoan that divides in red blood cell of mosquito as well as human (Amoudi, Alyousif et al. 2015). The most lethal form of malaria is caused by *P. falciparum* and *P. vivax* but in latter days *P. knowlesi* has emerged as severe to mankind also. It started infecting humans apart from its natural host i.e. long tailed macaques and rhesus macaques. So a detailed study of its genomic pattern is needed to understand the species in order to overcome from disease. This study uses available whole genome datasets for comparative genome analysis of *P. falciparum, P. vivax* and *P. knowlesi* in terms of their codon usage distribution. The genetic code i.e. codons, is a key that translates the information embedded in DNA as a nucleotide sequence to a functional protein as amino acid sequence. There are 64 codons out of which 61 are used for coding 20 amino acids and the rest are stop codon. So, more than one codon code for single amino acid referred as synonymous codon and their usage is different for each amino acid. This differential use is termed as codon usage bias which varies in both interspecies and intraspecies level (Grantham, Gautier et al. 1981). The difference in usage of synonymous codon modulates the gene expression level, mRNA stability, protein folding, translation efficiency and its accuracy (Plotkin and Kudla 2011). Codon usage bias is a result of compositional mutational bias and translational selection occurs at different level of genes. Many prokaryotes and fungi genome sequence attributed that codon usage is closely related to the intergenic GC content (Hershberg and Petrov 2010). In many prokaryotes (bacteria) and eukaryotes the codon and amino acid usage is highly influenced by the compositional GC mutational bias (Knight, Freeland et al. 2001). Translational selection also plays a vital role in shaping the codon usage pattern. Translationally optimal codons are used more preferentially than other synonymous codon. Translation selection depends upon the availability of abundant tRNA concentration mainly in the interaction between anticodon and wobbles position of codon. The translation rate also affects the folding of protein (Tuller, Waldman et al. 2010, Spencer 2012). The variation in synonymous codon usage may alter the polypeptide discloser from the ribosomes (Ran and Higgs 2010). The accuracy in protein folding is inversely correlated to the rate of translation (Yu, Dang et al. 2015). Rare codons are used to slow down the translation rate and allow the proper folding of functional proteins (Sachs and Malaney 2002, Shabalina, Spiridonov et al. 2013). *Plasmodium* species employ the replacement of preferred codon in protein low-complexity regions which slowdown the ribosomal speed for the proper assembly of protein secondary structure and protect the mis-folding of multidomain protein (Frugier, Bour et al. 2010). Variation in the DNA replication mechanism and repair system of both leading and lagging sequence creates the additional codon usage bias (Wald, Alroy et al. 2012). The codon usage bias can be determined by the variation in GC content, effective number of codon, codon adaptation index and various other factors. The GC content is closely associated with the evolution status of the organism (Muto and Osawa 1987, Behura and Severson 2013). Any similarities in the usage pattern determine the degree of biological relationship, environment adaptation and evolution among *Plasmodium* species (Yadav and Swati 2012), which can be used for the study of the molecular mechanism of translation, drug discovery, gene cloning and other biological functions among *Plasmodium* species.

## Materials and methods

### Genome Sequence Data

The complete coding sequences (CDSs) of *P. knowlesi, P. vivax and P. falciparum* were taken from PlasmoDB database (Aurrecoechea, Brestelli et al. 2009). Total number of CDSs analyzed: 5484 of *P. knowlesi,* 5610 of *P. vivax* and 5818 of *P. falciparum*.

### GC composition

The variation in the nucleotide composition of a genome can affect the synonymous codon choices for coding amino acid sequences. This correlation was firstly formulated by Sueoka in 1960s and suggest that it depend upon both mutational and selection pressure. Compositional bias at the DNA level affects both the synonymous and the non-synonymous sites in protein-coding genes (Rao, Wu et al. 2011), making the proteins to change their amino acid composition over evolutionary time (Grantham, Gautier et al. 1980). GC content is the percentage of either guanine or cytosine nucleotide bases of a DNA molecule.

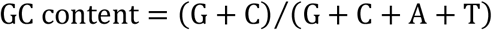

The GC content in intergenic region of a genome is correlated with the GC content of their genic sequences (Palidwor, Perkins et al. 2010). This influences the choice of codon to be either GC rich or AT rich. The third position of codon, GC3, allows the substitution of synonymous codons for coding amino acid (Yu and Li 2011). The codon is loosely bound to anticodon at the third position. This is a wobble position that allows the codon substitution without changing the amino acid sequence.

### Relative synonymous codon usage (RSCU)

The frequency of synonymous codons does not always reflect their true codon usage as the length of coding sequences differs in different genes in a genome. RSCU is a simple measure of non uniform usage of synonymous codons in a coding sequence, thus overcome the length effect. It is the number of times a particular codon is observed; relative to the number of times that codon would be observed for a uniform synonymous codon usage. The RSCU value of 1 indicates uniform usages of codons while less or more than 1 indicates their uneven usage.

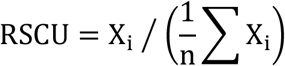

Where

X_i_ = occurrence of i^th^ synonymous codon of an n-fold degenerate amino acid.
n = total number of synonymous codons (1≤n≤6) for particular amino acid.

### Codon adaptation index(CAI)

CAI relays on the relative adaptiveness of the codon usage of highly expressed gene. The relative adaptiveness of each codon is the ratio of each codon, to that of the most abundant codon for the same amino acid (Sharp, Tuohy et al. 1986, Sharp and Devine 1989, Sharp, Emery et al. 2010). The CAI value for a gene is a measure of codon preference acting on any gene compared to the codon preferred in highly expressed proteins in a given genome.

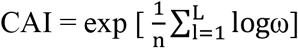

Where

L = number of codon in gene.
ω = relative adaptiveness of codon.

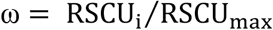

RSCU_i_ is the RSCU value for j^th^ codon for the i^th^ amino acid and RSCU_max_ is the RSCU value of the most frequently used codon for i^th^amino acid. The CAI of a gene ranges from 0 to 1; the higher value indicates the use of more preferred codons.

### Effective number of codon (ENc)

ENc is independent of gene length and amino acid composition and also a reliable estimator of codon usage bias in a gene. The value of ENc ranges from 20 to 61 where 61 values indicate random usage of codon while 20 values indicate biasness in the usage of codon (Wright 1990). The universal genetic code has five amino-acid family types (non-synonymous, 2-fold, 3-fold, 4-fold and 6-fold synonymous amino-acids). ENc is the arithmetic average of all non-zero homozygosity values (F) for each family type and the contributions from each of the synonymous families.

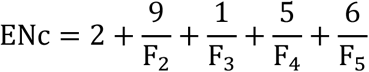

Where F is the codon homozygosity for amino acids having synonymous codons.

The ENc is closely related to GC3 composition of a gene. The relationship between ENc and GC3 can be approximated by the following equation:

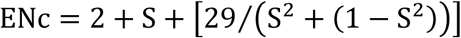

Where S is the frequency of GC3.

### Estimation of tRNA

The tRNA genes of all *Plasmodium* species were taken from GeneDB database (http://www.genedb.org) and their anticodons were detected by using tRNAscanSE (Lowe and Eddy 1997).

### General Average Hydrophobicity and Aromaticity

GRAVY (General Average Hydropathicity) value is the sum of the hydropathy values of all the amino acids in the gene product along the length of protein. The more negative the GRAVY value, the more hydrophilic the protein, while the more positive the GRAVY value, the more hydrophobic the protein (Kyte and Doolittle 1982). Aromo values denote the frequency of aromatic amino acids (Phe, Tyr, Trp) in the hypothetical translated gene product. The index and GRAVY value have been used to quantify the major COA trends in the amino acid composition of E. coli genes (Lobry and Gautier 1994, Jia, Liu et al. 2015).

### Correspondence analysis

Correspondence analysis (COA) is a multivariate statistical technique, used to investigate major trends in codon usage pattern. The technique is applied on different *Plasmodium* coding sequences using a CodonW program (written by J. Peden and available at http://www.molbiol.ox.ac.uk/cu). In COA, all genes were plotted in a 59-dimensional hyperspace corresponding to the RSCU values of 59 informative codons. Methionine, tryptophan and Stop codons were excluded from this study.

## Result and Discussion

### Base composition of *P. knowlesi* species indicate A/T bias

The *Plasmodium* species posses a similar, 14 number of chromosomes, but they differ in their genomic sizes. The genome size of *P. knowlesi* (23.46 Mb) falls in between other two human infecting *Plasmodium* species- *P. falciparum* (23.3 Mb) and *P. vivax* (27.01 Mb). The genomic GC percentage of *P. knowlesi, P. falciparum* and *P. vivax* are found to be 37.5 Mb, 19.41Mb and 42.3 Mb respectively. The coding region GC percentage of *P. knowlesi, P. falciparum* and *P. vivax* is 40.19 Mb, 23.78 Mb and 46.21Mb. The coding sequence GC percentage is higher than the genomic GC percentage with-in a species. This trend in also followed by studied *Plasmodium* species because coding parts of the genome are under higher GC selection pressure. There is an approximately equal contribution of all nucleotides in the coding part of *P. knowlesi* and *P. vivax*, indicates more or less random usage of synonymous codons. The genome of *P. falciparum* is considered as one of the most AT-rich genome, this is also reflected in their coding regions composition. The variation in GC composition in different regions of the genes lead to differential codon usage, and may be due to differential mutational pressure acting on different coding regions of a genome during the course of evolution.

Third codon position, also known as wobble position, is only moderately affected by natural selection and essentially immune from insertions and deletions, so it offers an opportunity to analyze long-term patterns of nucleotide substitutions. To evaluate in what extent *Plasmodium* exhibits GC-heterogeneity in their CDS’s, we have plotted the GC3 vs. number of coding sequence data as shown in Figure 1. In *P. knowlesi* GC3 ranges from 0.032 to 0.652, in *P. vivax* it ranges from 0.055 to 0.9 and in *P. falciparum* it ranges from 0 to 0.788. The GC3 distribution graph shows a wider distribution of genes, and a moderate use of GC at third codon position in *P. knowlesi* and *P. vivax* while the distribution of most of the genes of *P. falciparum* are biased towards low GC3 values (Figure 1). Thus, the role of genomic composition in making the choice of synonymous codons in *P. knowlesi* and *P. vivax* is very less compared to *P. falciparum*.

**Figure 1:**
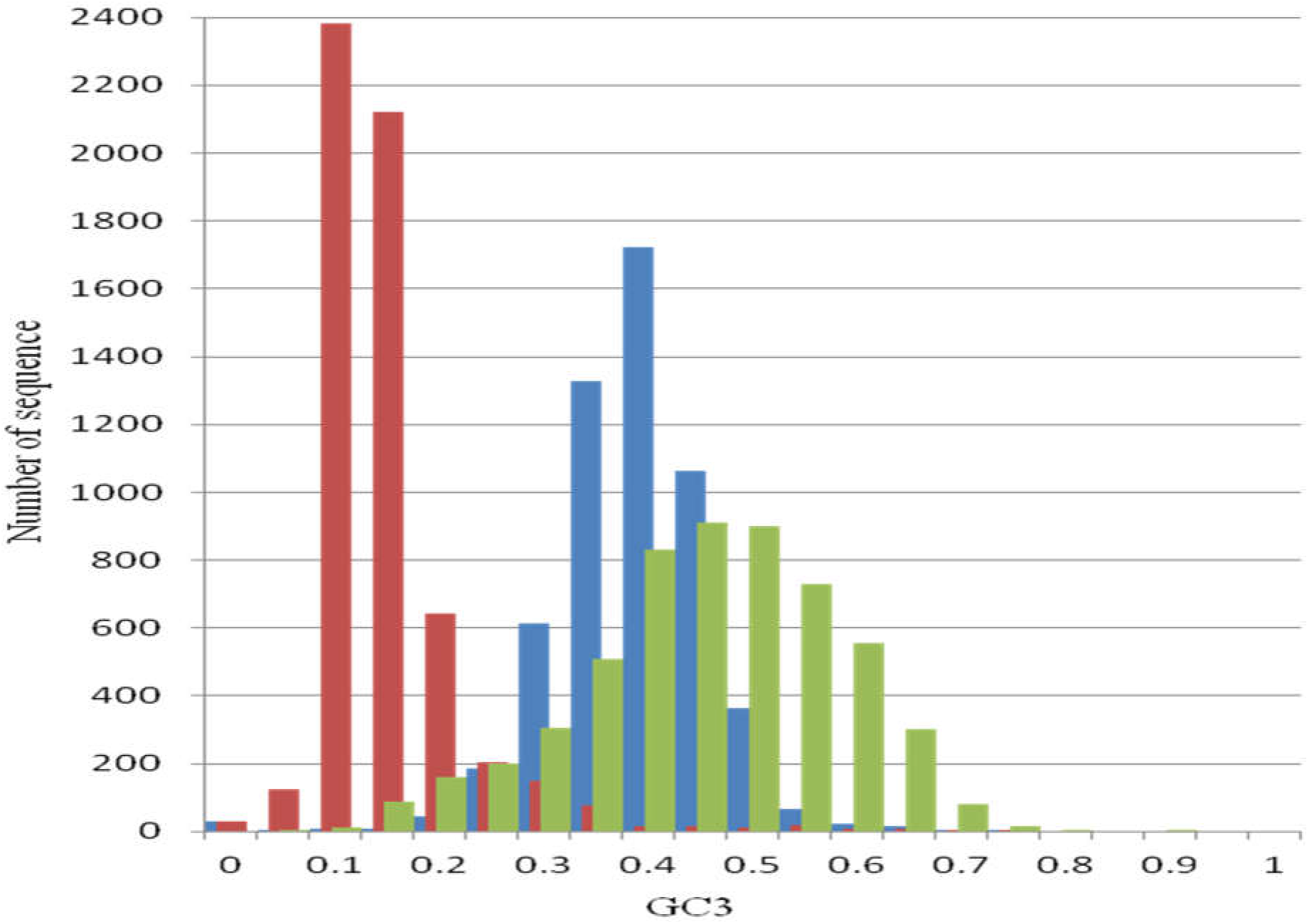
GC3 distribution in *P. knowlesi* (blue), *P. vivax* (Lang and Greenwood) and *P. falciparum* (red) coding sequences.

### Codon usage characteristics of *Plasmodium* species

Codon usage pattern of *Plasmodium* species varies in a noticeable manner between, and with-in the species. The relative synonymous codon usage (RSCU) is an important measure which is independent of amino acid usage, and may provide a better estimate of codon preferences. The RSCU distribution of *P. falciparum, P. vivax* and *P. knowlesi* genes are shown in Figure 2. From this, it is clear that the pattern of codon usage is relatively similar in *P. knowlesi* and *P. vivax.* Some codons are present in higher frequency for encoding a particular amino acid in a gene. These are called preferred codons. The present study has identified 18 preferred codons from the coding part of each *Plasmodium* species. The identified preferred codons of *P. knowlesi* use A/U bases at third codon position with an exception of G-ending Leu and Val, and C-ending Tyr amino acids. Higher use of A/U ending codons in *P. knowlesi* raises our concern about the role of genome composition in shaping their codon preferences. Usage of G/C ending codons will increase the overall GC bias, and this phenomenon is observed in the coding sequences of *P. vivax* where most of the codons are G/C ending except Ile, Lys and Phe amino acids. While all the preferred codons in *P. falciparum* coding sequences were found to be A/U ending. This A/U richness at the wobble position is supposed to be due to the A/U richness of their genome.

**Figure 2:**
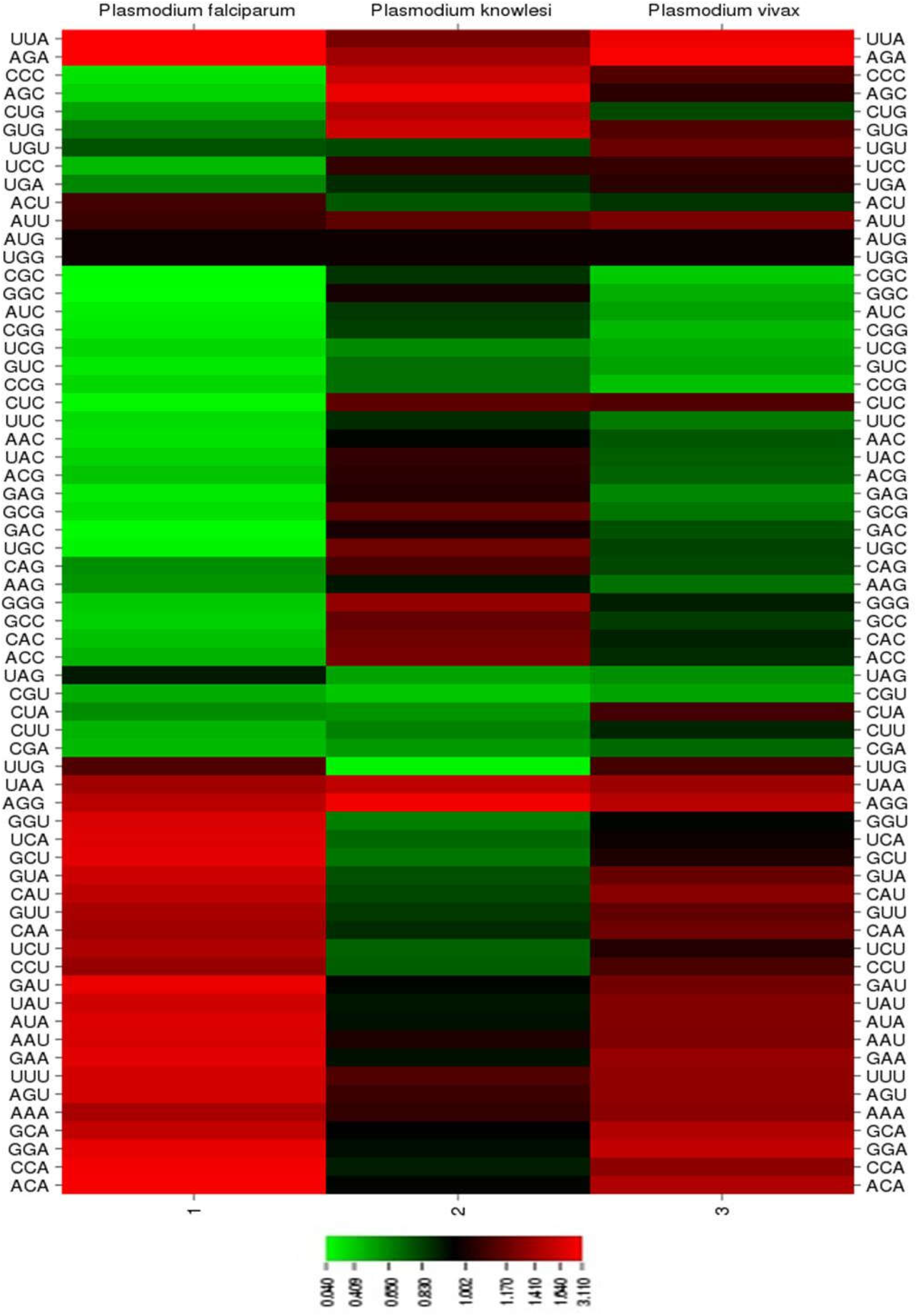
Codon usage pattern of CDS in *Plasmodium* species. Heat map was drawn in CIMminer using RSCU values of codons, by quantile binning method.

The pattern of codon usage bias bifurcates the *Plasmodium* species into two distinct groups in which one group comprises of *P. vivax* and *P. knowlesi,* and another group is of *P. falciparum* (Yadav and Swati 2012). This is not the case, while studying these species in the light of composition of preferred codons where *P. falciparum* and *P. knowlesi* were more A/U rich at the third codon position compared to their counterpart’s *P. vivax*.

The effective number of codons (ENc) is an advance measure, often used to assess the magnitude of codon bias of an individual gene and is independent of the gene length and frequency of amino acids. For a particular gene, ENc value is inversely proportional to their codon usage bias. This is used to evaluate the degree of codon usage bias in studied *Plasmodium* species.

The ENc distribution of protein coding genes of *P. knowlesi* and *P. vivax* is high compared to *P. falciparum*. This is clearly evident from average ENc calculation in *P. knowlesi* (54.25), *P. vivax* (53.72) and *P. falciparum* (39.06). Both, *P. knowlesi* and *P. vivax* shows more random usage of codons compared to that of *P. falciparum,* where codon preferences for large number of genes are more biased.

### Identification of Optimal Codons

Optimal codons are those codons that occur significantly more frequently in highly expressed genes compared to other genes (Lloyd and Sharp 1991). These codons are used frequently to increase the translation accuracy by preventing mis-folding errors (Gouy and Gautier 1982). The use of optimal codons in highly expressed genes permits more efficient use of ribosomes, and thus leads to a faster growth rate (Kudla, Murray et al. 2009). We have used ENc statistics for dividing the genes into highly and lowly expressed genes (Table 1). Top and bottom 5 % of genes were taken into account according to their ENc values. A statistical chi-squared test is performed on RSCU values of both the dataset; the codons that occurs frequently (*P*-value < 0.05) are considered as optimal codons.

**Table 1:** Codon usage table of highly and lowly expressed genes in *P. knowlesi, P. vivax* and *P. falciparum.*

| Amino acid | *P. knowlesi* |  |  |  | *P. vivax* |  |  |  | *P. falciparum* |  |  |  |
| --- | --- | --- | --- | --- | --- | --- | --- | --- | --- | --- | --- | --- |
|  | codon | RSCU<sup>a</sup> | RSCU<sup>b</sup> | tRNA | codon | RSCU<sup>a</sup> | RSCU<sup>b</sup> | tRNA | codon | RSCU<sup>a</sup> | RSCU<sup>b</sup> | tRNA |
| Phe | UUU* | 1.54 | 0.82 | 0 | UUU* | 1.51 | 1.02 | 0 | UUU* | 1.73 | 1.51 | 0 |
|  | UUC | 0.46 | 1.18 | 3 | UUC | 0.49 | 0.98 | 2 | UUC | 0.27 | 0.49 | 2 |
| Leu | UUA* | 2.85 | 0.37 | 2 | UUA* | 2.64 | 0.75 | 1 | UUA* | 4.53 | 1.66 | 1 |
|  | UUG | 0.95 | 1 | 2 | UUG | 0.93 | 1.28 | 1 | UUG | 0.5 | 1.69 | 1 |
|  | CUU* | 1.04 | 0.46 | 1 | CUU* | 1.04 | 0.78 | 1 | CUU | 0.5 | 0.52 | 1 |
|  | CUC | 0.25 | 1.58 | 0 | CUC | 0.37 | 1.11 | 0 | CUC | 0.08 | 0.27 | 0 |
|  | CUA | 0.65 | 0.53 | 2 | CUA* | 0.63 | 0.76 | 2 | CUA | 0.36 | 0.82 | 2 |
|  | CUG | 0.27 | 2.06 | 1 | CUG | 0.4 | 1.32 | 1 | CUG | 0.03 | 1.05 | 1 |
| Ile | AUU* | 1.34 | 0.98 | 1 | AUU* | 1.26 | 1.13 | 1 | AUU* | 1.27 | 0.82 | 1 |
|  | AUC | 0.33 | 1.35 | 1 | AUC | 0.32 | 0.96 | 0 | AUC | 0.15 | 0.35 | 0 |
|  | AUA* | 1.33 | 0.67 | 1 | AUA* | 1.43 | 0.91 | 1 | AUA | 1.58 | 1.84 | 1 |
| Met | AUG | 1 | 1 | 3 | AUG | 1 | 1 | 4 | AUG | 1 | 1 | 5 |
| Val | GUU* | 1.53 | 0.61 | 1 | GUU* | 1.38 | 0.89 | 1 | GUU* | 1.8 | 0.95 | 1 |
|  | GUC | 0.33 | 0.91 | 0 | GUC | 0.39 | 0.76 | 0 | GUC | 0.17 | 0.45 | 0 |
|  | GUA* | 1.64 | 0.46 | 2 | GUA* | 1.46 | 0.82 | 2 | GUA* | 1.79 | 1.38 | 3 |
|  | GUG | 0.5 | 2.03 | 1 | GUG | 0.77 | 1.54 | 1 | GUG | 0.24 | 1.22 | 1 |
| Tyr | UAU* | 1.53 | 0.5 | 0 | UAU* | 1.57 | 0.73 | 0 | UAU* | 1.86 | 1.34 | 0 |
|  | UAC | 0.47 | 1.5 | 2 | UAC | 0.43 | 1.27 | 2 | UAC | 0.14 | 0.66 | 2 |
| TER | UAA | 1.64 | 0.51 | 0 | UAA | 1.42 | 0.87 | 0 | UAA | 2.12 | 1.37 | 0 |
|  | UAG | 0.5 | 1.06 | 0 | UAG | 0.56 | 0.86 | 0 | UAG | 0.28 | 0.99 | 0 |
|  | UGA | 0.86 | 1.43 | 0 | UGA | 1.02 | 1.26 | 0 | UGA | 0.6 | 0.65 | 0 |
| His | CAU* | 1.55 | 0.5 | 0 | CAU* | 1.44 | 0.81 | 0 | CAU* | 1.8 | 1.23 | 0 |
|  | CAC | 0.45 | 1.5 | 1 | CAC | 0.56 | 1.19 | 2 | CAC | 0.2 | 0.77 | 2 |
| Gln | CAA* | 1.52 | 0.63 | 2 | CAA* | 1.39 | 0.92 | 2 | CAA* | 1.82 | 1 | 2 |
|  | CAG | 0.48 | 1.37 | 1 | CAG | 0.61 | 1.08 | 1 | CAG | 0.18 | 1 | 1 |
| Asn | AAU* | 1.57 | 0.68 | 0 | AAU* | 1.48 | 0.93 | 0 | AAU* | 1.79 | 1.37 | 0 |
|  | AAC | 0.43 | 1.32 | 2 | AAC | 0.52 | 1.07 | 2 | AAC | 0.21 | 0.63 | 1 |
| Lys | AAA* | 1.56 | 0.7 | 2 | AAA* | 1.49 | 0.99 | 2 | AAA* | 1.71 | 1.09 | 3 |
|  | AAG | 0.44 | 1.3 | 1 | AAG | 0.51 | 1.01 | 1 | AAG | 0.29 | 0.91 | 1 |
| Asp | GAU* | 1.65 | 0.69 | 0 | GAU* | 1.4 | 1.01 | 0 | GAU* | 1.83 | 1.38 | 0 |
|  | GAC | 0.35 | 1.31 | 2 | GAC | 0.6 | 0.99 | 2 | GAC | 0.17 | 0.62 | 1 |
| Glu | GAA* | 1.67 | 0.71 | 3 | GAA* | 1.6 | 1.09 | 2 | GAA* | 1.81 | 1.3 | 1 |
|  | GAG | 0.33 | 1.29 | 1 | GAG | 0.4 | 0.91 | 1 | GAG | 0.19 | 0.7 | 1 |
| Ser | UCU* | 1.44 | 0.41 | 1 | UCU* | 1.34 | 0.64 | 1 | UCU* | 1.63 | 0.92 | 1 |
|  | UCC | 0.57 | 1.29 | 0 | UCC | 0.86 | 1.32 | 0 | UCC | 0.37 | 0.78 | 0 |
|  | UCA* | 1.64 | 0.29 | 2 | UCA* | 1.34 | 0.6 | 2 | UCA* | 1.8 | 1.43 | 2 |
|  | UCG | 0.33 | 0.77 | 1 | UCG | 0.42 | 0.65 | 1 | UCG | 0.18 | 0.84 | 1 |
|  | AGU* | 1.5 | 0.86 | 0 | AGU* | 1.43 | 1.23 | 0 | AGU | 1.79 | 1.24 | 0 |
|  | AGC | 0.52 | 2.38 | 2 | AGC | 0.61 | 1.56 | 2 | AGC | 0.22 | 0.79 | 2 |
| Pro | CCU* | 1.58 | 0.41 | 1 | CCU* | 1.35 | 0.77 | 1 | CCU* | 1.64 | 0.83 | 1 |
|  | CCC | 0.58 | 2 | 0 | CCC | 0.8 | 1.51 | 0 | CCC | 0.19 | 0.6 | 0 |
|  | CCA* | 1.55 | 0.67 | 1 | CCA* | 1.56 | 1.01 | 1 | CCA* | 2.11 | 1.64 | 2 |
|  | CCG | 0.29 | 0.92 | 0 | CCG | 0.29 | 0.72 | 1 | CCG | 0.07 | 0.93 | 1 |
| Thr | ACU* | 1.33 | 0.42 | 1 | ACU* | 1.11 | 0.62 | 1 | ACU* | 1.24 | 0.77 | 1 |
|  | ACC | 0.45 | 1.66 | 0 | ACC | 0.62 | 1.29 | 0 | ACC | 0.42 | 0.66 | 0 |
|  | ACA* | 1.78 | 0.5 | 3 | ACA* | 1.72 | 1.07 | 1 | ACA* | 2.1 | 1.8 | 3 |
|  | ACG | 0.44 | 1.41 | 1 | ACG | 0.55 | 1.02 | 1 | ACG | 0.24 | 0.77 | 1 |
| Ala | GCU* | 1.34 | 0.52 | 1 | GCU* | 1.27 | 0.72 | 1 | GCU* | 1.97 | 0.94 | 1 |
|  | GCC | 0.37 | 1.38 | 0 | GCC | 0.52 | 1.22 | 0 | GCC | 0.29 | 0.67 | 0 |
|  | GCA* | 1.95 | 0.72 | 2 | GCA* | 1.76 | 1.09 | 1 | GCA* | 1.68 | 1.12 | 1 |
|  | GCG | 0.34 | 1.38 | 1 | GCG | 0.44 | 0.97 | 1 | GCG | 0.05 | 1.27 | 1 |
| Cys | UGU* | 1.42 | 0.42 | 0 | UGU* | 1.38 | 0.83 | 0 | UGU* | 1.86 | 1.23 | 0 |
|  | UGC | 0.58 | 1.58 | 1 | UGC | 0.62 | 1.17 | 1 | UGC | 0.14 | 0.77 | 1 |
| Trp | UGG | 1 | 1 | 1 | UGG | 1 | 1 | 1 | UGG | 1 | 1 | 1 |
| Arg | CGU* | 0.88 | 0.25 | 1 | CGU* | 0.59 | 0.5 | 1 | CGU* | 0.57 | 0.47 | 1 |
|  | CGC | 0.26 | 1.12 | 0 | CGC | 0.29 | 0.81 | 0 | CGC | 0.03 | 0.23 | 0 |
|  | CGA* | 0.72 | 0.52 | 2 | CGA* | 0.8 | 0.65 | 1 | CGA | 0.32 | 0.62 | 1 |
|  | CGG | 0.18 | 1.07 | 0 | CGG | 0.31 | 0.75 | 0 | CGG | 0.01 | 0.52 | 0 |
|  | AGA* | 2.94 | 0.89 | 2 | AGA* | 2.77 | 1.4 | 2 | AGA* | 4.61 | 1.75 | 3 |
|  | AGG | 1.02 | 2.16 | 1 | AGG | 1.24 | 1.9 | 1 | AGG | 0.46 | 2.41 | 1 |
| Gly | GGU* | 1.42 | 0.47 | 0 | GGU* | 1.13 | 0.81 | 0 | GGU* | 1.88 | 0.99 | 0 |
|  | GGC | 0.32 | 1.18 | 2 | GGC | 0.35 | 0.74 | 1 | GGC | 0.07 | 0.49 | 1 |
|  | GGA* | 1.82 | 0.76 | 1 | GGA* | 1.78 | 1.24 | 1 | GGA* | 1.82 | 1.52 | 2 |
|  | GGG | 0.43 | 1.58 | 0 | GGG | 0.73 | 1.21 | 0 | GGG | 0.22 | 1 | 0 |

N = codon count,
RSCU = Relative Synonymous codon usage,
RSCU<sup>a</sup> = RSCU of codon in highly expressed genes,
RSCU<sup>b</sup> = RSCU of codon in lowly expressed genes.
Note: Codon usage was compared using Chi squared contingency test to identify optimal codons that occur significantly more often (P<0.01) are indicated with \*.

The genome wide study reveals that only a select set of codons is optimized in each gene. The number of optimal codons used in *P. knowlesi* and *P. vivax* is more compared to *P. falciparum* and they are also unequal in distribution. The highly expressed gene of *P. knowlesi* has 29 optimal codon i.e. three for Ser (UCU, UCA, AGU), Arg (CGU, CGA, AGA), two for Leu (UUA, CUU), Pro (CCU, CCA), Ile (AUU, AUA), Val (GUU, GUA), Ala (GCU, GCA), Gly (GGU, GGA), Thr (ACU, ACA), one for His (CAU), Gln (CAA), Tyr (UAU), Phe (UUU), Cys (UGU), Lys (AAA), Asp (GAU), Glu (GAA), Asn (AAU) are estimated (Table 1). Highly expressed genes of *P. vivax* contain 30 optimal codons i.e. three for Ser (UCU, UCA, AGU), Arg (CGU, CGA, AGA), two for Leu (UUA, CUU, CUA), Pro (CCU, CCA), Ile (AUU, AUA), Val (GUU, GUA), Ala (GCU, GCA), Gly (GGU, GGA), Thr (ACU, ACA), one for His (CAU), Gln (CAA), Tyr (UAU), Phe (UUU), Cys (UGU), Lys (AAA), Asp (GAU), Glu (GAA), Asn (AAU) are estimated. While the highly expressed genes of *P. falciparum* show higher usage of 26 optimal codons i.e. Phe (UUU), Leu (UUA), Ile (AUU), Val (GUU, GUA), Tyr (UAU), His (CAU), Gln (CAA), Asn (AAU), Lys (AAA), Asp (GAU), Glu (GAA), Ser (UCU, UCA, AGU), Pro (CCU, CCA), Thr (ACU, ACA), Ala (GCU, GCA), Cys (UGU), Arg (CGU, AGA), Gly (GGU, GGA).

*P. falciparum* genome shows higher usage of A/T ended codons in their coding sequence, hence it is under high pressure of compositional mutational pressure while GC ended codon are the result of translational selection pressure acting on it (Bunnik, Chung et al. 2013). The comparatively GC rich species: *P. vivax* and *P. knowlesi*, show more use of A/T bases at wobble position of optimal codons indicates a prominent role of translational selection pressure acting at AT3 codons.

### Codon usage bias and tRNA copy number

The total copy number of tRNA in *P. knowlesi*, *P. vivax* and *P. falciparum* is 68, 67 and 57 respectively. Optimal codon usage highly recruits the most abundant tRNAs present in the organism having corresponding anticodon. Anticodon can encode for synonymous codons by wobble and supperwobble mechanism (Crick 1966). Superwobbling refers to the capability of an unmodified U in the wobble position of single tRNA species to base pair with all four nucleotides of a codon family (Rogalski, Karcher et al. 2008). In two-fold degenerate codons encoding amino acid i.e. Phenylalanine, Tyrosine, Histidine, Asparagine, Aspartate and cysteine tRNA set is present for less preferred codon than the preferred ones where the triplet with U at the third position is frequently used employing supperwobbling while in Glutamine, Lysine and Glutamate, more tRNAs set are present for preferred than non preferred codon for encoding polypeptide chain (Table 1). A supperwobbling between U:G base pairs are processed at a faster rate by ribosomes presumably because the wobble-G pairs more rapidly with the U codon than C (Higgs and Ran 2008). Amino acid with six fold degenerate codon, Arginine has more number of tRNA for preferred codons (AGA, CGA,CGT) and tRNA is absent for less preferred codon (CGC, CGG) in all the three species except AGG. For Leucine tRNAs are present for all preferred codons except CTA in *P. knowlesi* and CTT, CTA in *P. falciparum.* In Serine, supperwobbling occurs between AGC and AGT while optimal codon show more number of tRNA set.

The four fold degenerate codon encoding amino acids, in all the three species Valine, Proline, Threonine and alanine has more number of tRNA for preferred codon and tRNA is absent for less preferred codon expect CAC in Valine, CGG in Proline, CGT in Threonine and CGC in Alanine. In Glycine supperwobbling results in frequent usage of GGU.

Isoleucine has three fold degenerate codons where tRNA is present for preferred codons in all species as well as for less preferred codon TAT in *P. falciparum.*

The presence of tRNA for non-optimal codons of few amino acids in all the three species is to maintain the stability of proteins by avoiding the very strong as well as very weak codon: anticodon interactions. These extreme interactions increases the probability of mistranslation by ribosomes (Parmley and Huynen 2009). The use of alternating codons would slow down the translation process, required for correct folding of proteins.

### Codon usage adaptation in *Plasmodium* species

Codon adaptation index is an another method to assess the extent to which selection has been effective in molding the pattern of codon usage (Angov, Hillier et al.). CAI metrics are used to study the codon usage in highly and lowly expressed genes of studied *Plasmodium* species. It specifies the gene expression level under the assumption that translational selection pressure is available to optimize gene sequences. The average CAI value of coding region in *P. knowlesi* is 0.186 ranges from 0.045 to 0.622, *P. vivax* is 0.187 ranges from 0.049 to 0.622, and *P. falciparum* is 0.15 ranges from 0.045 to 0.556. The comparatively higher and wider range of CAI values in *P. knowlesi* and *P. vivax* indicates that their coding parts are under the weak selection pressure than *P. falciparum*. Thus, result in obvious similar usage of synonymous codons in highly expressed genes and rest of the genes in both the species (Prabha, Singh et al. 2017).

Correlation analysis between CAI and GC3 shows a positive correlation in *P. knowlesi, P. vivax* and *P. falciparum,* i.e. 0.338, 0.386 and 0.029 respectively (Table 2). The highly and lowly expressed genes shows higher use G:C at the third position of codon in *P. knowlesi* and *P. vivax* while *P. falciparum* shows higher usage of AT ended codons in coding sequences.

**Table 2:**
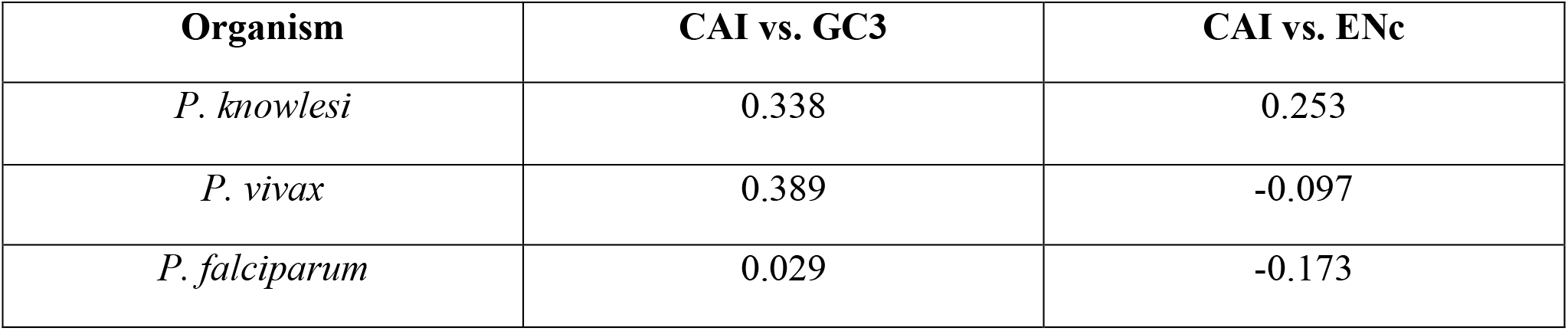
Correlation analysis between CAI, GC3 and ENc.

CAI shows weak but negative correlation with ENc i.e. −0.097 and −0.173 in *P. vivax* and *P. falciparum* respectively while has positive correlation in *P. knowlesi,* 0.253 (Table 2). The negative correlation shows that the gene expression influence the codon usage bias in the species but its effect is very negligible in *P. vivax* and the biasness occurs due to the confluence of genes having shorter length while in *P. falciparum* expression level influence the biasness in usage of synonymous codon. In *P. knowlesi* expression level plays a minor role encoding codon usage bias.

### Role of composition constraint and selection pressure

To better understand the relations between studies *Plasmodium* gene composition and codon usage bias, an ENC–GC3 scatter diagram was constructed. This method is usually used to estimate the role of important factors in shaping codon usage pattern. A common understanding about ENC–GC3 plot is that if a particular gene is subject to the GC compositional constraint to shape codon usage patterns, it will lie on a continuous curve, which represents a random codon usage. If a gene is subject to selection for translational optimal codons, it will lie considerably below the expected curve (Rudolph, Schmitt et al. 2016, Tyagi, Singh et al. 2016). The ENc value of each gene is plotted against its corresponding GC3 for *P. knowlesi*, *P. vivax* and *P. falciparum* (Figure 3). Although few genes in the three *Plasmodium* species were on the expected curve, several points were deviated from the solid line. The presence of genes in the middle of the curve on or below the solid line in *P. knowlesi* and *P. vivax* suggest more or less, a comparable role of both the mutation pressure and selection forces for shaping their codon usage pattern. While most of the genes of *P. falciparum* lies near the extreme left end of the normal curve, indicates the direct influence of GC3 composition. The expression-linked patterns of codon usage revealed that higher expression of genes was associated with lower GC3; lower ENc values in *P. falciparum*. The phenomenon is also evident by their correlation analysis where the values of correlation coefficient are higher (0.65) for *P. falciparum* compared to *P. knowlesi* (0.53) and *P. vivax* (0.15). The ENc-GC3 study will give a clear picture explaining the role of different forces in shaping their codon usage pattern, and suggest that the genes were not only subject to GC compositional constraints, but also natural selection in the studied three *Plasmodium* species. The codon usage bias in *P. falciparum* and *P. knowlesi*, more rely on GC compositional constraints apart from translation selection, in shaping their codon usage pattern than *P. vivax*.

**Figure 3:**
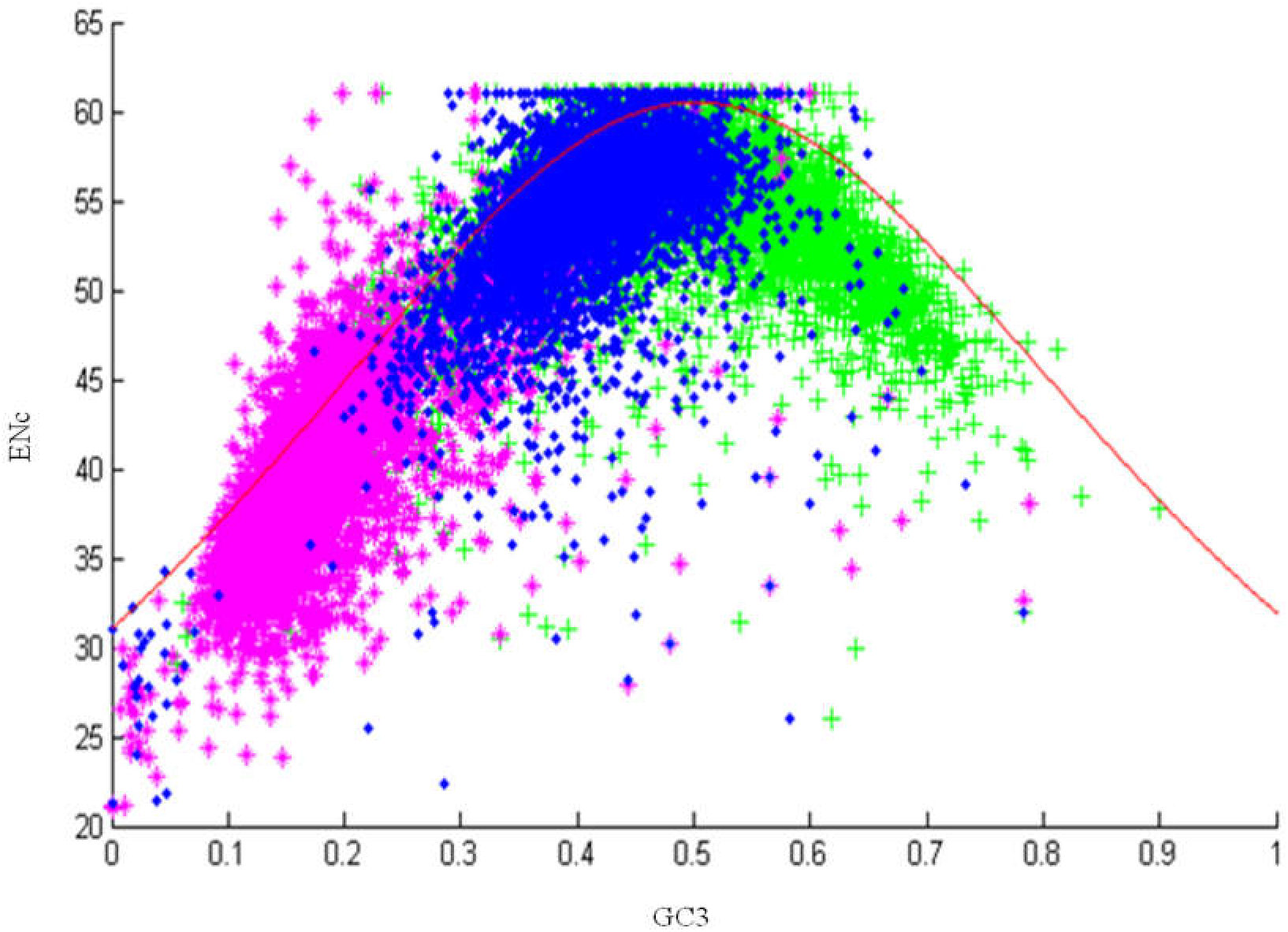
ENC versus GC3s plot. ENC denotes the effective number of codons of each gene, and GC3s denotes the G+C content on the third synonymous codon position of each gene of *P. knowlesi* (blue)*, P. vivax* (Lang and Greenwood) and *P. falciparum* (pink). The solid line (shown in red) indicates the expected ENC value if the codon bias is only due to GC3s.

### Correspondence analysis of the RSCU value

The codon usage variations among genes of *P. knowlesi, P. vivax* and *P. falciparum* are investigated by using correspondence analysis (COA) of their RSCU. A COA of the codon usage of the initial dataset composed of 5483 *P. knowlesi* chromosomal genes yielded a first axis that explained 9.4% of the total variation in this Codon Usage data. The variation explained by the second axis (5.5%) was approximately half of the variation explained by the first axis, with each subsequent axis explaining a decreasing amount of the variation (Fig. 4). The total variation explained by the two axes in other two *Plasmodium* species is shown in Table 3. The comparative COA result among the three *Plasmodium* species revealed a single major trend in codon usage, namely, that the first axis is the main contributor of the total variation compared to the next 3 axes, confirming that the primary axis is the main factor explaining codon usage in these genes (Table 3). The plot of first two axes, primarily, axis-1 and axis-2 for each gene is shown in Figure 5. The distance between genes on the plot is a reflection of their diversity in RSCU, with respect to the first two axes. Genes with similar codon usage are plotted as neighbors. Highly expressed genes preferentially using a subset of optimal codons (e.g. genes encoding glycolytic enzymes and or ribosomal proteins) are located on the left-hand side of the principal axis (Andersson and Sharp 1996b; Gouy and Gautier 1982; Lloyd and Sharp 1993; Sharp et al. 1988). Genes that one might predict to be moderately expressed, such as genes involved in amino-acid biosynthesis lie around the center of axis 1, while genes involved in regulation, lie at the other extreme of the principal axis (Bera, Virmani et al. 2017). The ordination of genes on the first two COA axes was examined for correlations with indices of codon usage and amino acid composition (e.g. GC3s, CAI, ENc, GRAVY, and Aromaticity). A summary of these correlations is presented in Table 4.

**Figure 4:**
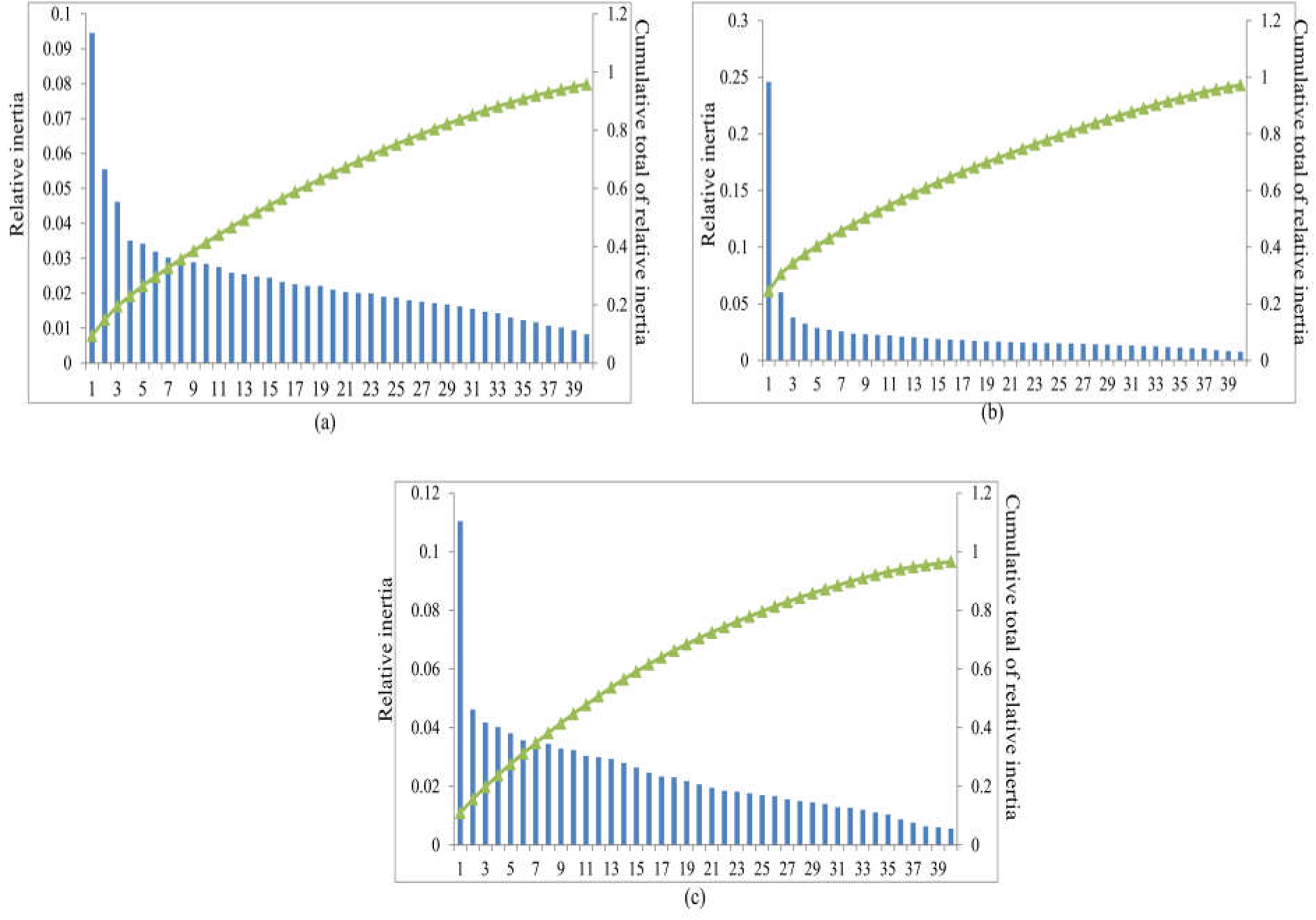
Each column represents a correspondence analysis factor of the codon usage of *P. knowlesi* (a)*, P. vivax* (b) and *P. falciparum* (c) ranked in decreasing order of the fraction of total variance or inertia in codon usage. The line represents the cumulative total of the inertia explained by the first 40 factors.

**Figure 5:**
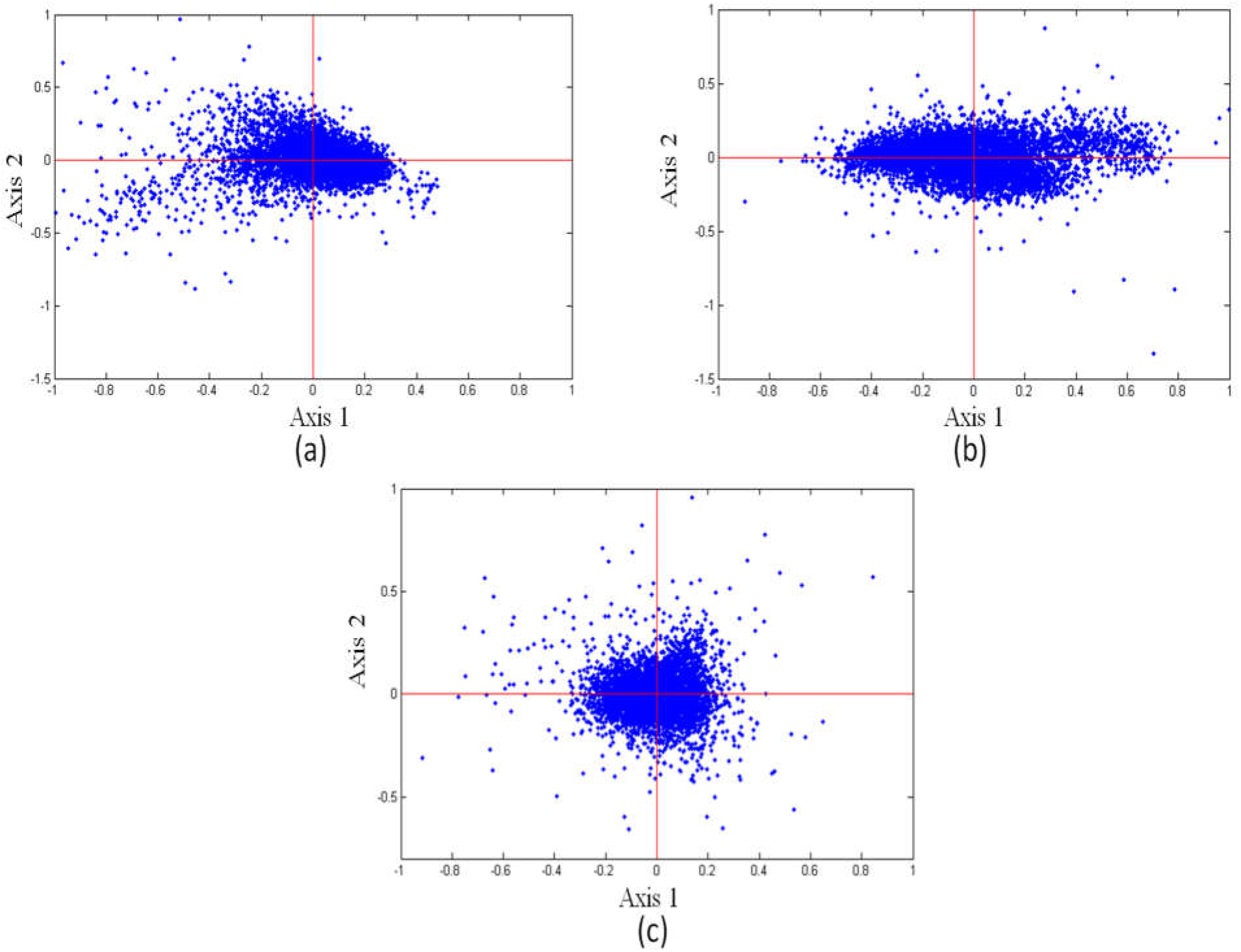
Correspondence analysis of codon usage variation among of *P. knowlesi* (a)*, P. vivax* (b) and *P. falciparum* (c) genes. Each gene is plotted using its coordinate on the first two axes produced by the analysis.

**Table 3:** Variation in codon usage along the axis generated by correspondence analysis.

| Axis | Organism |  |  |
| --- | --- | --- | --- |
|  | *P. knowlesi* | *P. vivax* | *P. falciparum* |
| 1 | 9.4% | 25% | 11% |
| 2 | 5.5% | 6% | 4.6% |

**Table 4:**
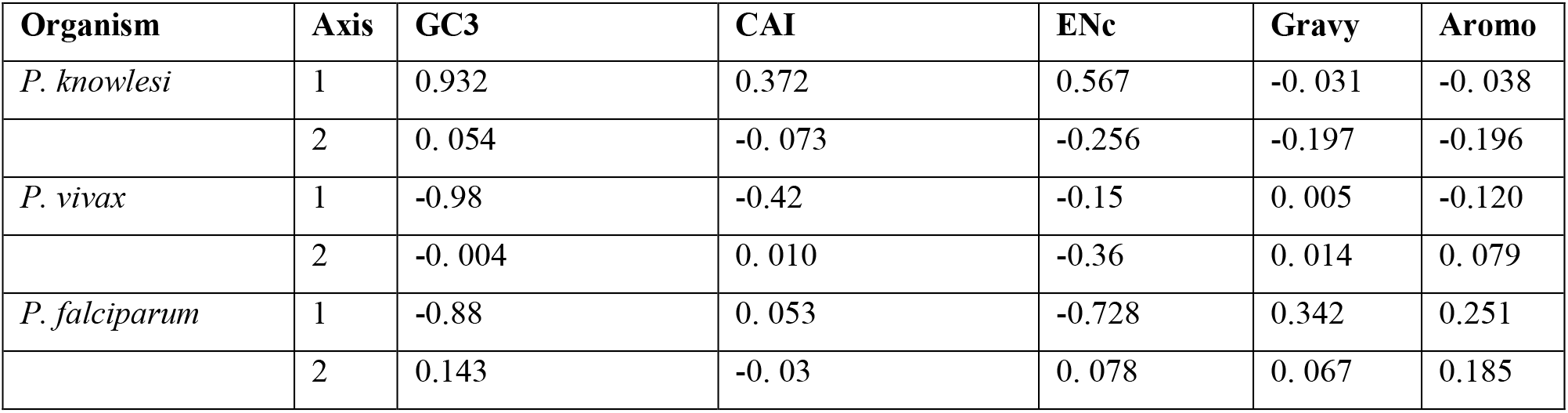
Correlation between codon usage indices and the Axis 1, Axis 2 ordination from a correspondence analysis.

| Organism | Axis | GC3 | CAI | ENc | Gravy | Aromo |
| --- | --- | --- | --- | --- | --- | --- |
| *P. knowlesi* | 1 | 0.932 | 0.372 | 0.567 | -0.031 | -0.038 |
|  | 2 | 0.054 | -0.073 | -0.256 | -0.197 | -0.196 |
| *P. vivax* | 1 | -0.98 | -0.42 | -0.15 | 0.005 | -0.120 |
|  | 2 | -0.004 | 0.010 | -0.36 | 0.014 | 0.079 |
| *P. falciparum* | 1 | -0.88 | 0.053 | -0.728 | 0.342 | 0.251 |
|  | 2 | 0.143 | -0.03 | 0.078 | 0.067 | 0.185 |

To identify the main contributor responsible for this variation in codon bias following correlation was calculated. In *P. knowlesi* Axis 1 shows significant correlation with GC3 (r=0.934, p<0.0001), CAI (r=0.372, p<0.0001) and ENc (0.567, p<0.0001) suggesting that both the composition mutation and translation selection pressure have an impact on the codon usage bias. Axis 1 is negative correlated with GC3, CAI and ENc in *P. vivax* indicating none of the selection pressure has a significant influence on the usage of synonymous codon thus the selection is random in *P. vivax.* In *P. falciparum* Axis 1 show negative correlation with GC3 and ENc while positive correlation with CAI indicating that the expression selection is the major factor whereas Axis 2 shows positive correlation with GC3 and ENc indicating that the composition biasness also influenced the non random usage of synonymous codon due to small variation of GC3 among the genes while the impact of translation selection pressure also have an effect on the codon usage bias. The results of codon usage analysis can also be influenced by the variation in amino acid composition which can be calculated by amino acid usage indices i.e., General average hydropathicity (GRAVY) and aromaticity (AROMO) (Rao, Wang et al. 2014). In *P. knowlesi* and *P. vivax* there is no significant corelation between the axes and the GRAVY and AROMO while in *P. falciparum* a positive correlation was observed indicating that effective selection of amino-acid for translational efficiency exists in this species (Xu, Cai et al. 2011).

## Conclusion

The codon usage pattern in the *Plasmodium* species is less conserved, changes occur in intraspecies genus level. The codon usage pattern of *P. knowlesi* was analyzed and compared with other human infect species*, P. falciparum* and *P. vivax.* Both the mutational composition and translation selection pressure play a significant role in creating the bias in the usage of synonymous codon in respective amino acid. The translational selection pressure play a significant role in all the three studied species while composition variation plays major role in *P. falciparum* that have low GC content creating higher biasness, in *P. knowlesi and P. vivax* mutation pressure also influences the usage but to a lesser extent (Chen and Cheng 1999, Chen, Lee et al. 2004). Correspondence analysis of all three species also indicates that the codon usage bias is the confluence of mutational bias and translational selection pressure. Thus the higher average ENC values, lower dependence on GC3 content and location of all genes on ENC-GC3 plot as in *P.knowlesi* shows similarity with that of *P. vivax*, thus shows more random usage of synonymous codons compared with that of *P.falciparum*. The CAI values in *P. knowlesi* and *P. vivax* which is very close to each other results in roughly equal nature of usage of codon for coding sequences as compared to that of *P. falciparum* which has very less CAI values showing higher non-random usage of synonymous codon. This pattern was also evidenced from various phylogenetic studies. The comparative codon usage study in *P. knowlesi* species gives insight to gene cloning experiments, gene expression and transfection studies of malaria species.

